# Slicing the genome of star-fruit (*Averrhoa carambola* L.)

**DOI:** 10.1101/851790

**Authors:** Yannan Fan, Sunil Kumar Sahu, Ting Yang, Weixue Mu, Jinpu Wei, Le Cheng, Jinlong Yang, Ranchang Mu, Jie Liu, Jianming Zhao, Yuxian Zhao, Xun Xu, Xin Liu, Huan Liu

## Abstract

The *Averrhoa carambola* is commonly known as star fruit because of its peculiar shape and its fruit is a rich source of minerals and vitamins. It is also used in traditional medicines in countries like India, China, the Philippines, and Brazil for treating various ailments such as fever, diarrhea, vomiting, and skin disease. Here we present the first draft genome of the Oxiladacea family with an assembled genome size of 470.51 Mb. In total, 24,726 protein-coding genes were identified and 16,490 genes were annotated using various well-known databases. Phylogenomic analysis confirmed the evolutionary position of the Oxiladacea family. Based on the gene functional annotations, we discovered the enzymes possibly involved in the biosynthesis of vitamin C, vitamin B_2_ and flavonoids pathways. Overall, being the first sequenced genome in the Oxiladacea family, the data provides an important resource for the nutritional, medicinal, and cultivational studies for this economically important star-fruit plant.

## Introduction

The star-fruit plant (*Averrhoa carambola* L.), a member of the Oxiladacea family, is a medium-size tree which is distinguished for its unique and attractive star shaped fruit. *A. carambola* is widely distributed around the world, especially in tropical countries, such as India, Malaysia, Indonesia, and the Philippines and is considered as an important species, and thus it is extensively cultivated in South-east Asia and Malaysia ^1,2^. Besides, it is also a popular fruit in the United States, Australia, and the South Pacific Islands market ^3^. Star-fruits have a special taste with slightly tart, acidic (smaller fruits) and sweet, mild flavor (large fruits). The star-fruit is known as a good source of various minerals and vitamins, and it is also rich in natural antioxidants such as vitamin C and gallic acid. Moreover, the presence of high amounts of fibers in fruits aids in absorbing glucose and retarding the glucose diffusion into the bloodstream and helps in controlling blood glucose concentration.

In addition to the food sources, the it is also utilized as herbs in India, Brazil, and Malaysia, and it is widely used in traditional Chinese Medicine preparations ^4^, as a remedy for fever, asthma, headache, and skin diseases ^5^. Several studies have demonstrated the presence of various phytochemicals such as saponins, flavonoids, alkaloids and tannins in the leaves, fruits, and roots of star-fruit plant ^6,7^, which are known to confer antioxidant and specific healing properties. The study of Cabrini *et al.,* ^5^ indicated that the ethanolic extract from *A. carambola* are highly useful in minimizing the symptoms of ear swelling (edema) and cellular migration in mice. The flavonoid compound (apigenin-6-C-β-fucopyranoside), which isolated from *A. carambola* leaves, showed the anti-hyperglycemic action in rats, and this might be a potential treatment and prevention of diabetes^8^. Moreover, the DMDD (2-dodecyl-6-methoxycyclohexa-2,5-diene-1,4-dione) extracted from the root of *A. carambola* exhibits potential benefit against obesity, insulin resistance, and memory deficits in Alzheimer’s disease ^9,10^.

Even though *A. carambola* play very significant roles in traditional medicine appalments, there are very limited studies at the genetic level on *A. carambola.,* mainly due to the lack of genome information. Therefore, filling this genomic gap will help the researchers to fully explore and understand this agriculturally important plant. As a part of 10KP project ^11,12^, in this study, the draft genome of *A. carambola* collected from Ruili botanical garden, Yunnan, China was assembled using advanced 10X genomics technique to further understand the evolution of Oxiladacea family. Furthermore, a fully annotated genome of *A. carambola* would serve as a foundation for the pharmaceutical applications and in the improvement of breeding strategies of the star-fruit plant.

## Results

### Genome assembly and evaluation

Based on the k-mer analysis, the total number of 35,655,285,391 kmers were used, with a peak coverage is 75. The *A. carambola* genome was estimated to be ~475 Mb in size (**Supplementary Fig. 1**). To perform the assembly, a total of 156 Gb clean reads were utilized by Supernova v2.1.1. The final assembly contained 69,402 scaffold sequences with N50 of 275.76 Kb and 78,313 contig sequences with N50 of 44.84 Kb for a total assembly size of 470.51 Mb (**Table 1**). Completeness assessment was performed using BUSCO (Bench-marking Universal Single-Copy Orthologs) version 3.0.1^13^ with Embryophyta odb9. The result showed that 1327 (92.20%) of the expected 1440 conserved plant orthologs were detected as complete (**Supplementary Table 1**).

**Table 1.** Statistics of genome assembly.

|  | Contig |  | Scaffold |  |
| --- | --- | --- | --- | --- |
|  | Size(bp) | Number | Size(bp) | Number |
| **N90\*** | 5,420 | 12,457 | 7,033 | 3,988 |
| **N80** | 13,875 | 7,548 | 39,210 | 608 |
| **N70** | 23,325 | 5,165 | 34,6109 | 131 |
| **N60** | 33,460 | 3,619 | 1,307,770 | 60 |
| **N50** | 44,841 | 2,503 | 2,757,598 | 35 |
| **Longest** | 717,770 | - | 14,768,062 | - |
| **Total Size** | 431,262,337 | - | 470,508,511 | - |
| **Total number**<br>**(≥2kb)** | - | 18,820 | - | 10,777 |
| **Total number**<br>**(≥100bp)** | - | 78,313 | - | 69,402 |

\*: Nxx length is the maximum length L such that xx% of all nucleotides lie in contigs (or scaffolds) of size at least L.

### Genome annotation

A total of 68.15% of the assembled *A. carambola* genome was composed of repetitive elements (**Supplementary Table 2**). Among these repetitive sequences, the LTRs were the most abundant, accounting for 61.64% of the genome. DNA class repeat elements represented 4.19% of the genome; LINE and SINE classes encoded 0.28% and 0.016% of the assembled genome, respectively. For the gene prediction, we combined homology- and *de novo*- based approaches and obtained a non-redundant set of 24,726 gene models with 4.11 exon per gene on average. The gene length was 3,457 bp on average, while the average exon and intron lengths were 215 bp and 827 bp, respectively. The gene model statistics data compared with other seven homology species are shown in Supplementary Fig 3. Functions were assigned to 16,490 (66.69%) genes. These protein-coding genes were then subjected for further exploration against KEGG, NR and COGs protein sequence databases ^14^, in addition to SwissProt, and TrEMBL ^15^, and then InterProScan ^16^ was lastly used to identify domains and motifs (**Supplementary Table 3, Supplementary Fig. 3**). Non-coding RNA genes in the assembled genome were also annotated. We predicted 759 tRNA, 1,341 rRNA, 90 microRNA (miRNA) and 2,039 small nuclear RNA (snRNA) genes in the assembled genome (**Supplementary Table 4**).

### Gene family analysis and phylogenetic tree

We performed *A. carambola* gene family analysis using OrthoMCL software ^17^, with protein and nucleotide sequences from *A. carambola* and ten other plant species (*A. thaliana, C. sinensis, F. sylvatica, G. max, K. fedtschenkoi, M. domestica, P. granatum, P. trichocarp, T. cacao, V. vinifera*) based on an all-versus-all BLASTP alignment with an E-value cutoff of 1e-05. The predicted 24,726 protein-coding genes in *A. carambola* were assigned to 9,731 gene families consisting of 15,301 genes, while 9,425 were not organized into groups which were specious unique in *A. carambola* (**Supplementary Table 5**). In total, 163 single-copy orthologs corresponding to the eleven species were extracted from the clusters and were used to construct the phylogenetic tree. The constructed tree topology supported the APG IV ^18^ system that Oxalidales (*A. carambola*) and Malpighiales (*P. trichocarp*) belong to the same cluster Rosids. Based on the phylogenetic tree, *A. carambola* was estimated to separate from *P. trichocarp, V. vinifera* and *K. fedtschenkoi* approximately 94.5, 110.2 and 126.3 Mya, respectively (**Supplementary Fig. 4**). Since *A. carambola* is the first species with whole-genome sequenced and assembled in the Oxalidaceae family, it is crucial to identify the evolutionary position of this family.

We also analyzed the expansion and contraction of the gene families between species using CAFÉ. The result showed that 888 gene families were substantially expanded and 15,724 gene families were contracted in *A. carambola* (**Fig. 1**), in total, 2,916 and 6,057 genes of *A. carambola* were identified from expanded and contracted families, respectively. which the contraction was about 17 times more than expansion.

**Figure 1.**
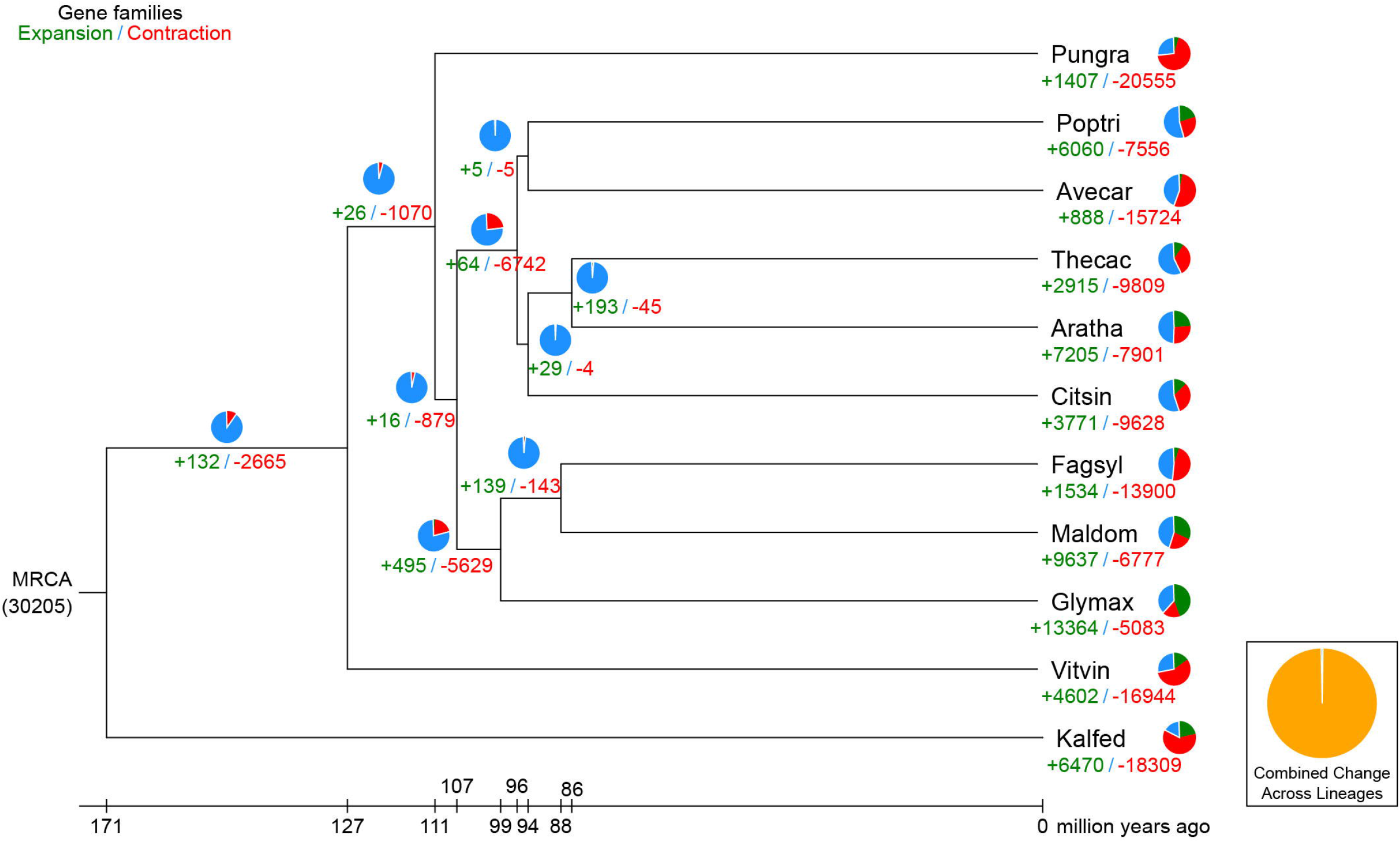
Phylogenetic tree showing the evolution of gene family sizes. Pungra: *P. granatum;* Poptri: *P. trichocarpa;* Avecar: *A. carambola;* Thecac: *T. cacao;* Aratha: *A. thaliana;* Citsin: *C. sinensis;* Fagsyl: *F. sylvatica;* Maldom: *M. domestica;* Glymax: *G. max*; Vitvin: *V. vinifera*; Kalfed: *K. fedtschenkoi.*

Later, the Gene Ontology (GO) and KEGG functional enrichment analyses were performed for all expansion gene families. The KEGG pathway enrichment analysis results are shown in Table 2 and the GO enrichment are listed in Supplementary Table 6. In a previous study, researchers had isolated the flavonoids from the fresh fruit of *A. carambola,* which are known to reduce the harmful inflammation ^19^. In our study, the flavonoid biosynthesis pathway was found to be significantly enriched in the expanded families. Terpenoids are yet another important type of compound which has been isolated from star fruit^20^, which has been proven to exhibit anti-inflammatory activities. *A. carambola* likely synthesize terpenoids via the Diterpenoid biosynthesis pathway.

**Table 2.** Enriched KEGG pathways of unique genes of *A. carambola* with expansion.

| Pathway ID | KEGG description | Adjusted<br>P-value<br>($\leq 0.05$) | Number<br>of genes | Total genes<br>in the<br>pathway |
| --- | --- | --- | --- | --- |
| map03020 | RNA polymerase | 3.51E-09 | 30 | 137 |
| map00904 | Diterpenoid biosynthesis | 1.00E-05 | 18 | 63 |
| map00240 | Pyrimidine metabolism | 3.16E-05 | 34 | 215 |
| map01100 | Metabolic pathways | 7.17E-05 | 234 | 2621 |
| map00901 | Indole alkaloid biosynthesis | 7.17E-05 | 15 | 55 |
| map00230 | Purine metabolism | 7.17E-05 | 36 | 244 |
| map00565 | Ether lipid metabolism | 0.00014092 | 14 | 52 |
| map01110 | Biosynthesis of secondary metabolites | 0.00038039 | 142 | 1478 |
| map02010 | ABC transporters | 0.00080624 | 20 | 119 |
| map00902 | Monoterpenoid biosynthesis | 0.00116462 | 9 | 29 |
| map00460 | Cyanoamino acid metabolism | 0.00270665 | 18 | 110 |
| map00941 | Flavonoid biosynthesis | 0.03063434 | 17 | 120 |
| map00190 | Oxidative phosphorylation | 0.0354555 | 19 | 142 |
| map00940 | Phenylpropanoid biosynthesis | 0.0354555 | 32 | 280 |
| map00062 | Fatty acid elongation in mitochondria | 0.0354555 | 7 | 32 |
| map00563 | Glycosylphosphatidylinositol(GPI)-anchor<br>biosynthesis | 0.03937265 | 8 | 41 |
| map00040 | Pentose and glucuronate interconversions | 0.0426558 | 21 | 167 |

### Identification of vitamin C, vitamin B_2_, and flavonoids pathway

The star-fruit is an excellent source of various minerals and vitamins, especially for natural antioxidants such as L-ascorbic acid (Vitamin C) and flavonoids ^1,19^. Through the ortholog search in the KEGG pathway, we identified the enzymes which are potentially involved in the vitamin C, vitamin B_2_ and flavonoid synthesis pathway in *A. carambola.*

In a previous report, a major component of plant ascorbate was reported to synthesize through the l-galactose pathway ^21^, in which GDP-d-mannose is converted to l-ascorbate by four successive intermediates, as summarized in Fig. 2A. Laing *et al.* ^22^ reported the identification of l-galactose guanylyltransferase encoding homologous genes from *Arabidopsis* and kiwifruit which catalyzes the conversion of GDP-l-galactose to l-galactose-1-P In this study, five necessary enzymes - GalDH, GalLDH, GGalPP, GalPP, and GME, were identified, which are involved in the vitamin C pathway, suggesting the possibility of ascorbic acid synthesis in the star-fruit (Table 3). On the other hand, we also identified the possible enzymes involved in riboflavin (vitamin B_2_) biosynthesis pathway in the star-fruit (Fig. 2B, Table 3). Through the catalyzation by RIB3, RIB4, and RIB5, the D-Ribulose 5-phosphate compound can finally produce the riboflavin.

**Figure 2.**
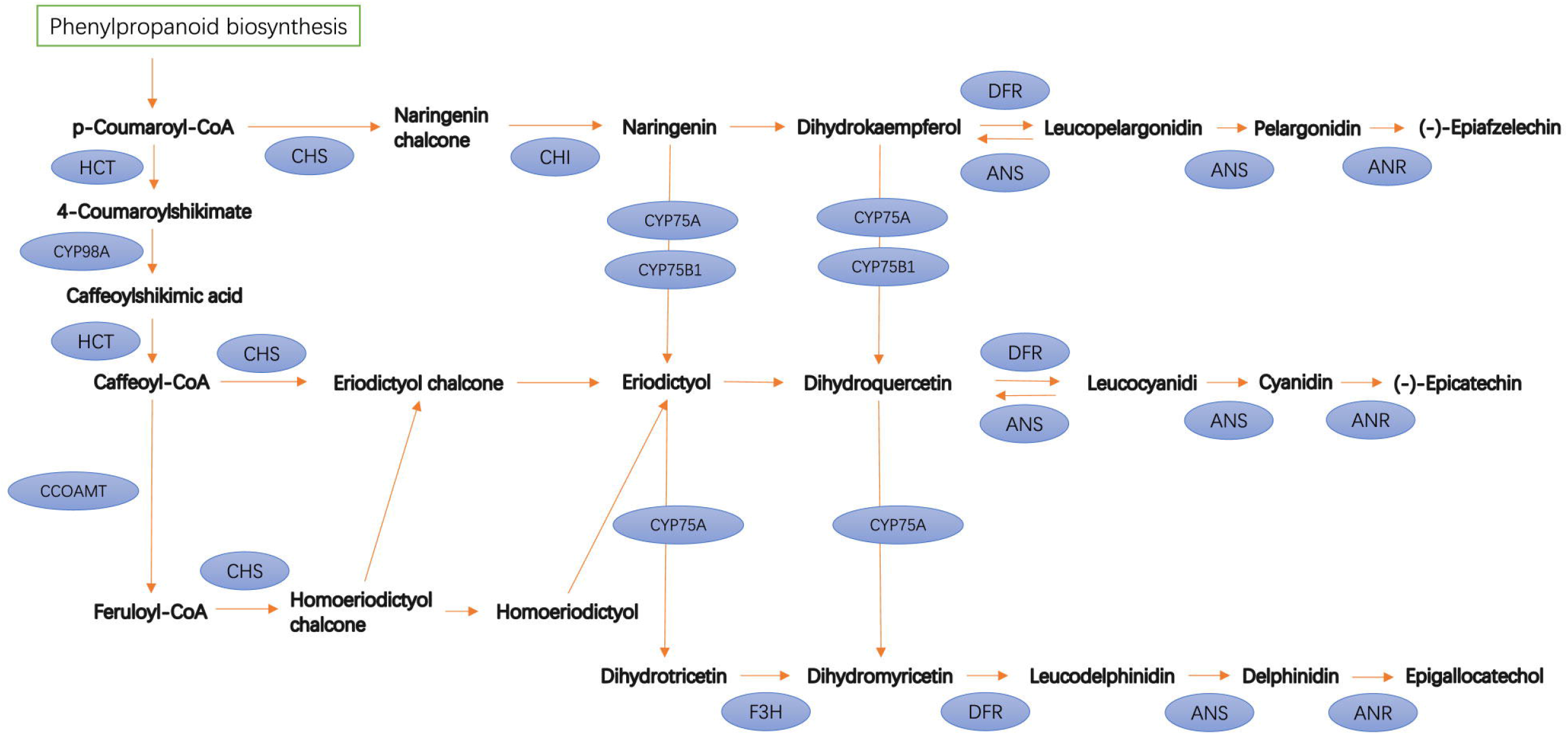
Identification of genes involved in the (a) Vitamin C pathway and (b) Vitamin B_2_ pathway. Genes identified as being involved in two pathways shown in the blue circle. GalDH: L-galactose dehydrogenase; GalLDH: L-galactono-1,4-lactone dehydrogenase; GGalPP: GDP-L-galactose phosphorylase; GalPP: L-galactose-1-phosphate phosphatase; GME: GDP-D-mannose-3’,5’-epimerase; FLAD1: FMN adenylyltransferase; RIB3: 3,4-dihydroxy 2-butanone 4-phosphate synthase; RIB4: 6,7-dimethyl-8-ribityllumazine synthase; RIB5: riboflavin synthase; RFK: riboflavin 5’-phosphotransferase.

**Table 3.** List of genes involved in vitamin C and Vitamin B_2_ pathway.

| pathway | Enzyme | Description | Copy numbers | Gene ID | Protein (AA) |
| --- | --- | --- | --- | --- | --- |
| Vitamin C pathway | GalDH | L-galactose dehydrogenase | 4 | Aca.sc093668.g0.5 | 332 |
|  |  |  |  | Aca.sc000054.g1.6 | 334 |
|  |  |  |  | Aca.sc000055.g24.2 | 351 |
|  |  |  |  | Aca.sc000055.g24.3 | 594 |
|  | GalLDH | L-galactono-1,4-lactone dehydrogenase | 5 | Aca.sc000059.g19.35 | 601 |
|  |  |  |  | Aca.sc000229.g0.29 | 603 |
|  |  |  |  | Aca.sc000078.g43.5 | 598 |
|  |  |  |  | Aca.sc100684.g0.3 | 99 |
|  |  |  |  | Aca.sc102574.g0.3 | 99 |
|  | GGalPP | GDP-L-galactose phosphorylase | 3 | Aca.sc150475.g0.5 | 275 |
|  |  |  |  | Aca.sc150564.g0.4 | 288 |
|  |  |  |  | Aca.sc098705.g0.4 | 410 |
|  | GalPP | L-galactose-1-phosphate phosphatase | 2 | Aca.sc006151.g1 | 181 |
|  |  |  |  | Aca.sc000246.g13.57 | 360 |
|  | GME | GDP-D-mannose-3',5'-epimerase | 2 | Aca.sc096116.g0.5 | 383 |
|  |  |  |  | Aca.sc000061.g13.42 | 383 |
| Vitamin B<sub>2</sub> pathway | RIB3 | 3,4-dihydroxy 2-butanone 4-phosphate synthase | 3 | Aca.sc000063.g38.15 | 545 |
|  |  |  |  | Aca.sc000056.g65.27 | 572 |
|  |  |  |  | Aca.sc000071.g11.34 | 590 |
|  | RIB4 | 6,7-dimethyl-8-ribityllumazine synthase | 1 | Aca.sc023103.g76.2 | 185 |
|  | RIB5 | riboflavin synthase | 1 | Aca.sc000036.g2.16 | 343 |
|  | RFK | riboflavin 5'-phosphotransferase | 1 | Aca.sc000058.g77.42 | 539 |
|  | FLAD1 | FMN adenylyltransferase | 1 | Aca.sc000058.g25.58 | 520 |

In the *A. carambola* gene family analysis, the KEGG pathway enrichment analysis for the expanded gene families revealed that 17 genes participates in flavonoid synthesis pathway (P-value = 0.03, Table 3). The previous studies proved that flavonoids can be isolated from *A. carambola* and other plants from the Oxalidaceae family ^1,9,19,23–26^. Here, we identified 11 enzymes that could be potentially involved in the flavonoid biosynthesis pathway (Fig. 3, Supplementary Table 7). The two enzymes in the pathway which contain 23 and 21 homolog genes are HCT (shikimate O-hydroxycinnamoyltransferase) and CHS (chalcone synthase), respectively. Among the end-point products, apigenin, cyanidin, epicatechin, and quercetin has been extracted from the leaves, fruits or barks of *A. carambola* in previous studies ^6,27–29^.

**Figure 3.**
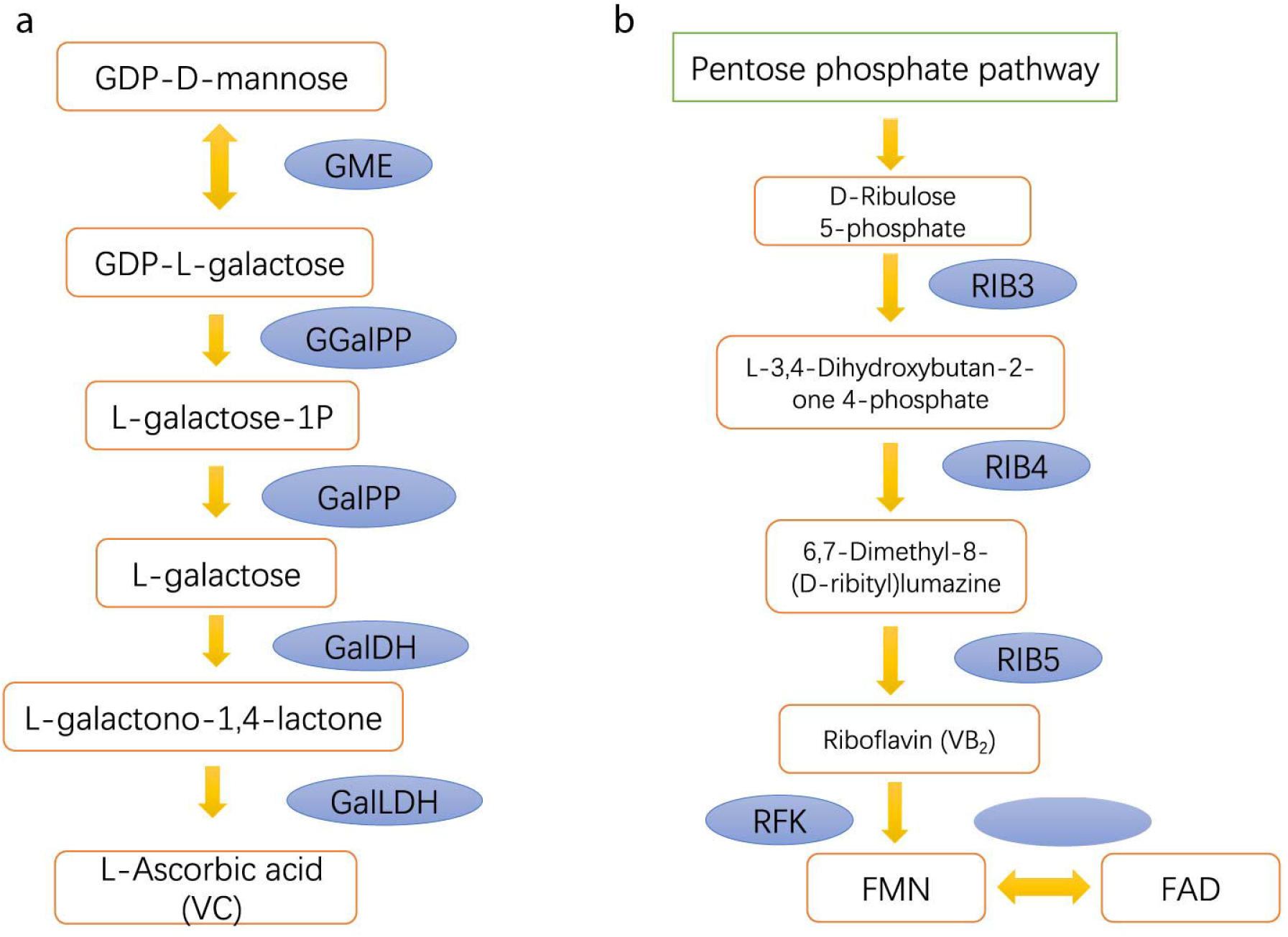
Identification of genes involved in the flavonoid biosynthesis pathway. Genes identified as being involved in two pathways shown in the blue circle. ANR: anthocyanidin reductase; ANS: leucoanthocyanidin dioxygenase; CCOAMT: caffeoyl-CoA O-methyltransferase; CHI: chalcone isomerase; CHS: chalcone synthase; CYP75A: flavonoid 3’,5’-hydroxylase; CYP75B1: flavonoid 3’-monooxygenase; CYP98A: coumaroylquinate 3’-monooxygenase; DFR: flavanone 4-reductase; F3H: naringenin 3-dioxygenase; HCT: shikimate O-hydroxycinnamoyltransferase.

## Discussion

This study presents the first draft genome in the Oxalidaceae family with relatively high-quality genome assembly. The sequenced species *A. carambola* (star fruit) is a widely cultivated and utilized as an edible fruit and serves as an important source of minerals, vitamins, and phytomedicinal properties. The genome size was assembled to be 470.51 Mb with scaffold N50 of 275.76 Kb. Since we cannot compare the genome size with other species in this family, we have found a similar genome size, which is 434.29 Mb of *Populus trichocarpa* and 350.62 Mb of *Ricinus communis* in the closest order Malpighiales. However, the genome of the chromosome level will be further required to better understand the diploid character of this species.

There are 24,726 gene models that have been identified from *A. carambola.* The gene number is relatively smaller than what has been reported in *A. thaliana, P. trichocarpa,* or *T. cacao.* The length distribution of exons of prediction star fruit proteins was consistent with other species, although its intron and CDS length tended to be shorter than other species in the comparison (**Supplementary Fig. 2**). The proportion of predicted genes that contain InterPro functional domain is 52.3%, and that can be aligned with NCBI nr database (66.4%) are the highest among all databases. It is likely that *A. carambola* is so far the only species which has been assembled in the Oxalidaceae family; there might be some evolutionary unique genes in this family remaining to be annotated.

Later we performed gene family analysis together with the other ten species and identified the significant expansion of 888 gene families contains 2,916 unique genes in *A. carambola.* These genes have significantly enriched (P-value <= 0.05) within 28 GO classes, including 18 biological processes, 2 cellular component, and 8 molecular functions (**Supplementary Table 6**). The DNA binding biological process contained the most genes (60) within the expanded families. Within the significantly enriched biological processes, defense response to fungus might can be related to the anti-microbial property of star fruits in the previous studies^1^. On the other hand, we found that oxidoreductase activity is enriched in the molecular function GO class, which could be one of the potential reasons for the antioxidant activity in star fruits. Moreover, within the enriched KEGG pathway, we identified the enzymes involved in three metabolic pathways – vitamin C, vitamin B_2_, and flavonoid pathway. Although the potential functional enzymes have been identified from the genome, those functional pathways should be further verified by the experimental studies in the future.

In the summary, it can be expected that this draft genome will facilitate in understanding the formation of specific and important traits in the star-fruit plant, such as the discovery of biosynthesis pathways of pharmacologically active metabolites, and in the improvement of breeding of strategies of the star-fruit plant.

## Materials and methods

### Plant materials and sequencing

The leaf samples of *A. carambola* were collected from the Ruili Botanical Garden, Yunnan, China. The genomic DNA was extracted by using CTAB (cetyltriethylammonium bromide) method ^30^. The 10X Genome sequencing was performed to obtain high-quality reads. The high-molecular-weight (HMW) DNA was loaded onto a Chromium Controller chip with 10X Chromium reagents and gel beads, and rest of the procedures were carried out as per the manufacturer’s protocol ^31^. Then, BGISEQ-500 platform to produce 2 × 100 bp paired-end sequences, generating a total of 183 Gb raw data. The raw reads were filtered using SOAPfilter v2.2 with the following parameters “-q 33 -i 600 -p -l -f -z -g 1 -M 2 -Q 20”. After filtering low-quality reads, around 157 Gb of clean data (high-quality reads >Q35) remained for the next step.

### Estimation of *A. carambola* genome size

All the 157 Gb clean reads obtained from the BGISEQ-500 platform were subjected to the 17 kmer frequency distribution analysis with Jellyfish ^32^ using the parameters “-k 17 −t 24”. The frequency graph was drawn, and the *A. carambola* genome size was calculated using the formula: genome size = k-mer_Number/Peak_Depth.

### *De novo* genome assembly

The Linked-Read data were assembled using the Supernova v2.1.1 software using the “--localcores=24 --localmem=350 --max reads 280000000” parameter. In order to fill the scaffold gaps, GapCloser version 1.12 ^33^ was used with the parameters “-t 12 −l 155”. Finally, the total assembled length of *A. carambola* genome was 470.51 Mb, with a scaffold N50 of 275.76 Kb and contig N50 of 44.84 Kb, respectively.

### Repeat annotation

For the transposable element annotation, RepeatMasker v3.3.0 ^34^ and RepeatProteinMasker v3.3.0 ^34^ were used against Repbase v16.10 ^35^ to identify known repeats in the *A. carambola* genome. Tandem repeats were identified using software Tandem Repeats Finder v4.07b ^36^. *De novo* repeat identification was conducted using RepeatModeler v1.0.5 ^37^ and LTR_FINDER v1.05 ^38^ programs, followed by RepeatMasker v3.3.0 ^34^ to obtain the final results.

### Gene prediction

Prior to the gene prediction analysis, we masked the repetitive regions of the genome. The MAKER-P v2.31 ^39^ was utilized to predict the protein-coding genes based on homologous, and *de novo* prediction evidence. For homologous-based prediction, protein sequences of *Theobroma cacao, Prunus persica, Prunus mume, Prunus avium, Populus trichocarpa, Populus euphratica,* and *Arabidopsis thaliana* were aligned against *A. carambola* genome using BLAT ^40^. Then, the gene structure was predicted using GeneWise ^41^. In order to optimize different *ab initio* gene predictors, we constructed a series of training set for *de novo* prediction evidence. Complete gene models by homologous evidence were picked for training in Augustus tool ^42^. Genemark-ES v4.21 ^43^ was self-trained using the default criteria. The first round of MAKER-P was also run using the default parameters on the basis of above evidences, with the exception for “protein2genome” which was set to “1”, yielding only protein-supported gene models. SNAP ^44^ was then trained with these gene models. Default parameters were used to run the second and the final rounds of MAKER-P, generating the final gene models.

### Functional annotation

The predicted gene models were functionally annotated by aligning their protein sequences against the Kyoto Encyclopedia of Genes and Genomes (KEGG) ^45^, the Clusters of Orthologous Groups (COG) ^14^, SwissProt ^15^, TrEMBL, and the National Center for Biotechnology Information (NCBI) non-redundant (NR) protein databases with BLASTP (E-value <= 1.0e-0.5). Protein motifs and domains were identified by comparing the sequences against various domain databases, including PFAM, PRINTS, PANTHER, ProDom, PROSITE, and SMART using InterProScan v5.21 ^16^. For ncRNA annotation, tRNA genes were identified by tRNAscan-SE v1.23 ^46^. For the identification of rRNA genes, we aligned the assembled data against the rRNA sequences of *A. thaliana* using BLASTN (E-value <= 1.0e-0.5). The miRNAs and snRNAs were predicted by using INFERNAL ^47^ software against the Rfam database ^48^.

### Gene family construction and phylogenetic analysis

For gene family analysis, the OrthoMCL ^17^ software was utilized to construct orthologous gene families on all the protein-coding genes of *A. carambola* and 12 sequenced plant species (*B. napus, G. raimondii, H. brasiliensis, J. curcas, L. usitatissimum, M. esculenta, P. euphratica, P. trichocarpa, P. mume, R. communis, T. cacao, V. vinifera).* Before OrthoMCL, the BLASTP was used to find similar matches from different species with an E-value cutoff of 1.0e-05. The composition of the OrthoMCL clusters was used to calculated the total number of gene families. Orthogroups that were single copy in all species analyzed were selected and aligned using MAFFT v7.310 ^49^. Each gene tree was constructed by RAxML v8.2.4 ^50^ with GTRGAMMA model. To construct the species phylogenetic tree, a coalescent-based method by ASTRAL v4.10.4 ^51^, with 100 replicates of multi-locus bootstrapping ^52^ were used.

The divergence time between *A. carambola* and other species were estimated using MCMCTREE ^50^ with the default parameter. The expansion and contraction of gene family numbers were predicted using the CAFÉ ^53^ by employing the phylogenetic tree and gene family statistics.

## Data availability

The data sets generated and analyzed during the current study are available in the CNGB Nucleotide Sequence Archive (CNSA: https://db.cngb.org/cnsa). The raw sequencing data is under ID CNR0066625, and assembly data is under ID CNA0002506. All other data generated or analyzed during this study are included in this published article and its supplementary information files.

## Acknowledgments

This work was supported by funding from the Shenzhen Municipal Government of China (grants JCYJ20170817145512476 and JCYJ20160510141910129), the Guangdong Provincial Key Laboratory of Genome Read and Write (grant 2017B030301011), and the NMPA Key Laboratory for Rapid Testing Technology of Drugs.

## Contributions

R.-c.M., J.L., J.-m.Z., T.Y., Y.-x.Z. and W.-x.M. collected the samples; W.-x.M and S.K.S. conceived and conducted the experiments; Y.-n.F. and T.Y. analyzed the results; Y.-n.F. and S.K.S wrote the manuscript.

