## Supplemental tables and figures for "Slicing the genome of star-fruit (*Averrhoa carambola* L.)"

**Supplementary Table 1.** Completeness evaluation of genome assembly using BUSCO database for *A. carambola*

| BUSCO | Number | Percentage |
|---|---|---|
| **Complete BUSCOs** | 1327 | 92.2% |
| **Complete single copy** | 1290 | 89.6% |
| **Complete duplicated** | 37 | 2.6% |
| **Fragmented** | 39 | 2.7% |
| **Missing** | 74 | 5.1% |

**Supplementary Table 2.** Categories of TEs predicted in the *A. carambola* genome.

| | Repbase TEs | | TE protiens | | De novo | | Combined TEs* | |
|---|---|---|---|---|---|---|---|---|
| Type | Length (Mb) | % in genome | Length (Mb) | % in genome | Length (Mb) | % in genome | Length (Mb) | % in genome |
| DNA | 6.37 | 1.35 | 2.03 | 0.43 | 11.29 | 2.40 | 19.69 | 4.19 |
| LINE | 0.64 | 0.14 | 0.03 | 0.004 | 0.67 | 0.14 | 1.34 | 0.28 |
| SINE | 0.47 | 0.01 | 0 | 0 | 0.03 | 0.06 | 0.07 | 0.016 |
| LTR | 62.53 | 13.29 | 29.36 | 6.24 | 198.14 | 42.11 | 290.03 | 61.64 |
| Other | 5.08 | 0.001 | 0 | 0 | 0 | 0 | 0.005 | 0.001 |
| Unknown | 0 | 0 | 0 | 0 | 14.88 | 3.16 | 14.88 | 3.16 |
| Total* | 69.15 | 14.70 | 30.15 | 6.41 | 221.35 | 47.05 | 320.64 | 68.15 |

*: the total number of TEs was identified by combining all repeats identified through different methods. As there are some overlaps between different methods, the combined number of TEs is less than the sum of repeats identified by all methods.

**Supplementary Table 3.** Gene annotation in the *A. carambola* genome.

| | **Number** | **Percent(%)** |
|---|---|---|
| **Total** | 24,726 | 100% |
| **Annotated** | 16,490 | 66.69% |
| **NR** | 16,418 | 66.40% |
| **KEGG** | 12,146 | 49.12% |
| **COG** | 6,170 | 24.95% |
| **SwissProt** | 12,937 | 52.32% |
| **InterPro** | 12,932 | 52.30% |
| **Unannotated** | 8,236 | 33.31% |

**Supplementary Table 4.** Non-coding RNA genes in the *A. carambola* genome.

| Type | | Copy(w) | Average length(bp) | Total length(bp) | % of genome |
|---|---|---|---|---|---|
| miRNA | | 90 | 130.68 | 11761 | 0.0025 |
| tRNA | | 759 | 75.46 | 57274 | 0.012173 |
| rRNA | rRNA | 1341 | 143.01 | 191777 | 0.04076 |
| | 18S | 583 | 206.28 | 120259 | 0.025559 |
| | 28S | 514 | 92.79 | 47692 | 0.010136 |
| | 5.8S | 104 | 96.08 | 9992 | 0.002124 |
| | 5S | 140 | 98.81 | 13834 | 0.00294 |
| snRNA | snRNA | 2039 | 107.60 | 219387 | 0.046628 |
| | CD-box | 1919 | 106.00 | 203422 | 0.043234 |
| | HACA-box | 34 | 121.76 | 4140 | 0.00088 |
| | splicing | 86 | 137.5 | 11825 | 0.002513 |

**Supplementary Table 5. Ortholog analysis of predicted genes in *A. carambola* with those annotated in other twelve plant species**

| Species | Genes number | Genes in families | Unclustered genes | Family number | Unique families | Average genes per family |
|---|---|---|---|---|---|---|
| *A. carambola* | 23946 | 15301 | 9425 | 9731 | 643 | 1.38 |
| *A. thaliana* | 48306 | 44857 | 3449 | 14405 | 1845 | 3.11 |
| *C. sinensis* | 35456 | 33813 | 1643 | 14494 | 655 | 2.33 |
| *F. sylvatica* | 31728 | 27722 | 4006 | 13192 | 806 | 2.1 |
| *G. max* | 88647 | 78854 | 9793 | 16626 | 2681 | 4.74 |
| *K. fedtschenkoi* | 45190 | 39422 | 5768 | 14224 | 1619 | 2.77 |
| *M. domestica* | 59695 | 54005 | 5690 | 15679 | 2037 | 3.44 |
| *P .granatum* | 29127 | 23129 | 5998 | 13039 | 833 | 1.77 |
| *P. trichocarpa* | 45468 | 37319 | 8149 | 15519 | 1137 | 2.4 |
| *T. cacao* | 30734 | 29609 | 1125 | 14528 | 328 | 2.04 |
| *V. vinifera* | 40694 | 38824 | 1870 | 14646 | 724 | 2.65 |

**Supplementary Table 6. Enriched GO terms of genes in families with expansion.**

| GO ID | GO Term | GO Class | Adjusted P-value (<= 0.05) | Num.of genes |
|---|---|---|---|---|
| GO:0009112 | nucleobase metabolic process | BP | 1.13E-05 | 6 |
| GO:0008610 | lipid biosynthetic process | BP | 0.00768689 | 14 |
| GO:0050832 | defense response to fungus | BP | 0.00949146 | 3 |
| GO:0016998 | cell wall macromolecule catabolic process | BP | 0.0112318 | 4 |
| GO:0006022 | aminoglycan metabolic process | BP | 0.0112318 | 4 |
| GO:0006040 | amino sugar metabolic process | BP | 0.0112318 | 4 |
| GO:0000226 | microtubule cytoskeleton organization | BP | 0.02281434 | 4 |
| GO:0043085 | positive regulation of catalytic activity | BP | 0.02440192 | 3 |
| GO:0042157 | lipoprotein metabolic process | BP | 0.02465928 | 4 |
| GO:0006631 | fatty acid metabolic process | BP | 0.02763653 | 8 |
| GO:0032787 | monocarboxylic acid metabolic process | BP | 0.03039913 | 8 |
| GO:1901136 | carbohydrate derivative catabolic process | BP | 0.03613591 | 4 |
| GO:0072527 | pyrimidine-containing compound metabolic process | BP | 0.04185329 | 4 |
| GO:0006664 | glycolipid metabolic process | BP | 0.04185329 | 4 |
| GO:0046467 | membrane lipid biosynthetic process | BP | 0.04185329 | 4 |
| GO:0016053 | organic acid biosynthetic process | BP | 0.04185329 | 12 |
| GO:0046394 | carboxylic acid biosynthetic process | BP | 0.04185329 | 12 |
| GO:0042545 | cell wall modification | BP | 0.04684068 | 6 |
| GO:0005815 | microtubule organizing center | CC | 0.02281434 | 4 |
| GO:0005783 | endoplasmic reticulum | CC | 0.03089248 | 6 |
| GO:0050661 | NADP binding | MF | 0.00027827 | 9 |
| GO:0003677 | DNA binding | MF | 0.00072907 | 60 |
| GO:0016709 | oxidoreductase activity, acting on paired donors, with incorporation or reduction of molecular oxygen, NAD(P)H as one donor, and incorporation of one atom of oxygen | MF | 0.00264374 | 7 |
| GO:0043531 | ADP binding | MF | 0.00368156 | 14 |
| GO:0016843 | amine-lyase activity | MF | 0.0112318 | 4 |
| GO:0005315 | inorganic phosphate transmembrane transporter activity | MF | 0.02440192 | 2 |
| GO:0016663 | oxidoreductase activity, acting on other nitrogenous compounds as donors, oxygen as acceptor | MF | 0.02440192 | 2 |
| GO:0016790 | thiolester hydrolase activity | MF | 0.02465928 | 4 |

**Supplementary Table 7. List of genes involved in Flavonoid biosynthesis pathway**

| Enzyme | Description | Copy number | Gene name | Protein (AA) |
|---|---|---|---|---|
| HCT | shikimate O-hydroxycinnamoyltransferase | 23 | Aca.sc06029.g3 | 444 |
| | | | Aca.sc096404.g1 | 443 |
| | | | Aca.sc000036.g82 | 438 |
| | | | Aca.sc091789.g1 | 101 |
| | | | Aca.sc000054.g1 | 438 |
| | | | Aca.sc000038.g7 | 168 |
| | | | Aca.sc089800.g0.4 | 356 |
| | | | Aca.sc088407.g0.25 | 970 |
| | | | Aca.sc000036.g40 | 114 |
| | | | Aca.sc097067.g0.13 | 447 |
| | | | Aca.sc089799.g1 | 440 |
| | | | Aca.sc000054.g80 | 133 |
| | | | Aca.sc000063.g80 | 451 |
| | | | Aca.sc824068.g1 | 428 |
| | | | Aca.sc231962.g2 | 314 |
| | | | Aca.sc000036.g40 | 202 |
| | | | Aca.sc000061.g10 | 201 |
| | | | Aca.sc096735.g1 | 133 |
| | | | Aca.sc098445.g1 | 299 |
| | | | Aca.sc061321.g1 | 374 |
| | | | Aca.sc000125.g5 | 466 |
| | | | Aca.sc000054.g81 | 453 |
| | | | Aca.sc000054.g80 | 453 |
| CYP98A | coumaroylquinate 3'-monooxygenase | 1 | Aca.sc000073.g35 | 503 |
| CCOAMT | caffeoyl-CoA O-methyltransferase | 6 | Aca.sc000059.g38 | 247 |
| | | | Aca.sc006197.g1 | 154 |
| | | | Aca.sc000289.g0.18 | 234 |
| | | | Aca.sc000289.g3 | 187 |
| | | | Aca.sc096535.g0.2 | 202 |
| | | | Aca.sc000063.g0.48 | 309 |
| CHS | chalcone synthase | 21 | Aca.sc000062.g4 | 768 |
| | | | Aca.sc0000181.g2 | 394 |
| | | | Aca.sc152842.g0.22 | 392 |
| | | | Aca.sc152841.g0.21 | 392 |
| | | | Aca.sc000075.g15 | 392 |
| | | | Aca.sc000231.g4 | 392 |
| | | | Aca.sc000075.g16 | 390 |
| | | | Aca.sc000054.g2.5 | 397 |
| | | | Aca.sc000054.g2.1 | 390 |

| | | | | |
|---|---|---|---|---|
| | | | Aca.sc000213.g8 | 387 |
| | | | Aca.sc091597.g0.3 | 398 |
| | | | Aca.sc000213.g7.8 | 387 |
| | | | Aca.sc091519.g0.4 | 292 |
| | | | Aca.sc103007.g6 | 354 |
| | | | Aca.sc000062.g5.4 | 992 |
| | | | Aca.sc118539.g0.2 | 263 |
| | | | Aca.sc000054.g30 | 396 |
| | | | Aca.sc096624.g1 | 255 |
| | | | Aca.sc000080.g3.4 | 391 |
| | | | Aca.sc000246.g21 | 383 |
| | | | Aca.sc000078.g10.4 | 1174 |
| CHI | chalcone isomerase | 3 | Aca.sc000058.g3.6 | 624 |
| | | | Aca.sc000081.g12.3 | 629 |
| | | | Aca.sc000134.g0.68 | 185 |
| ANS | leucoanthocyanidin dioxygenase | 5 | Aca.sc097807.g1.58 | 353 |
| | | | Aca.sc097807.g0.36 | 274 |
| | | | Aca.sc096691.g0.4 | 436 |
| | | | Aca.sc000219.g19 | 362 |
| | | | Aca.sc000058.g56 | 369 |
| F3H | naringenin 3-dioxygenase | 7 | Aca.sc000078.g45.7 | 342 |
| | | | Aca.sc000058.g44 | 352 |
| | | | Aca.sc000111.g9.32 | 366 |
| | | | Aca.sc000111.g9.5 | 67 |
| | | | Aca.sc095606.g0.4 | 129 |
| | | | Aca.sc000111.g9.33 | 157 |
| | | | Aca.sc093304.g1 | 141 |
| CYP75B1 | flavonoid 3'-monooxygenase | 1 | Aca.sc000073.g29 | 956 |
| CYP75A | flavonoid 3',5'-hydroxylase | 3 | Aca.sc098656.g1.6 | 511 |
| | | | Aca.sc097345.g1.3 | 511 |
| | | | Aca.sc087790.g1 | 171 |
| DFR | flavanone 4-reductase | 6 | Aca.sc000054.g64 | 340 |
| | | | Aca.sc000069.g13 | 121 |
| | | | Aca.sc000246.g38 | 335 |
| | | | Aca.sc110869.g0.4 | 342 |
| | | | Aca.sc000069.g6.6 | 162 |
| | | | Aca.sc000078.g36 | 348 |
| ANR | anthocyanidin reductase | 6 | Aca.sc000069.g6.68 | 162 |
| | | | Aca.sc000054.g85 | 641 |
| | | | Aca.sc000078.g36.3 | 348 |
| | | | Aca.sc000069.g13.6 | 121 |
| | | | Aca.sc110869.g0.41 | 342 |
| | | | Aca.sc000057.g13 | 571 |

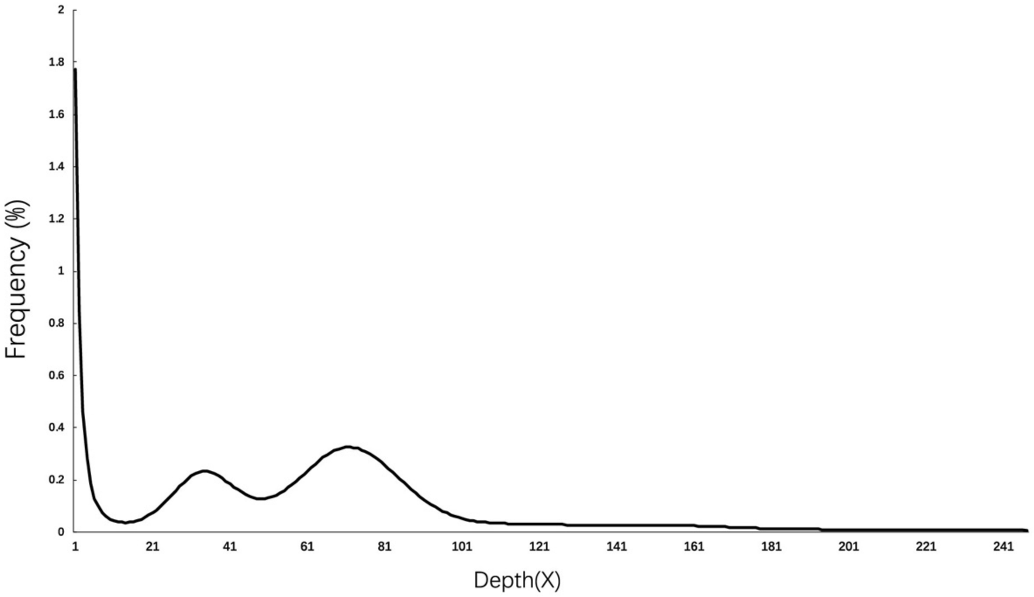


**Supplementary Figure 1. The 17-mer depth distribution of *A. carambola.*** The X-axis is depth; the Y-axis represents the frequency. According to the distribution, we estimate that the genome size of *A. carambola* is approximately 475.40 Mb.

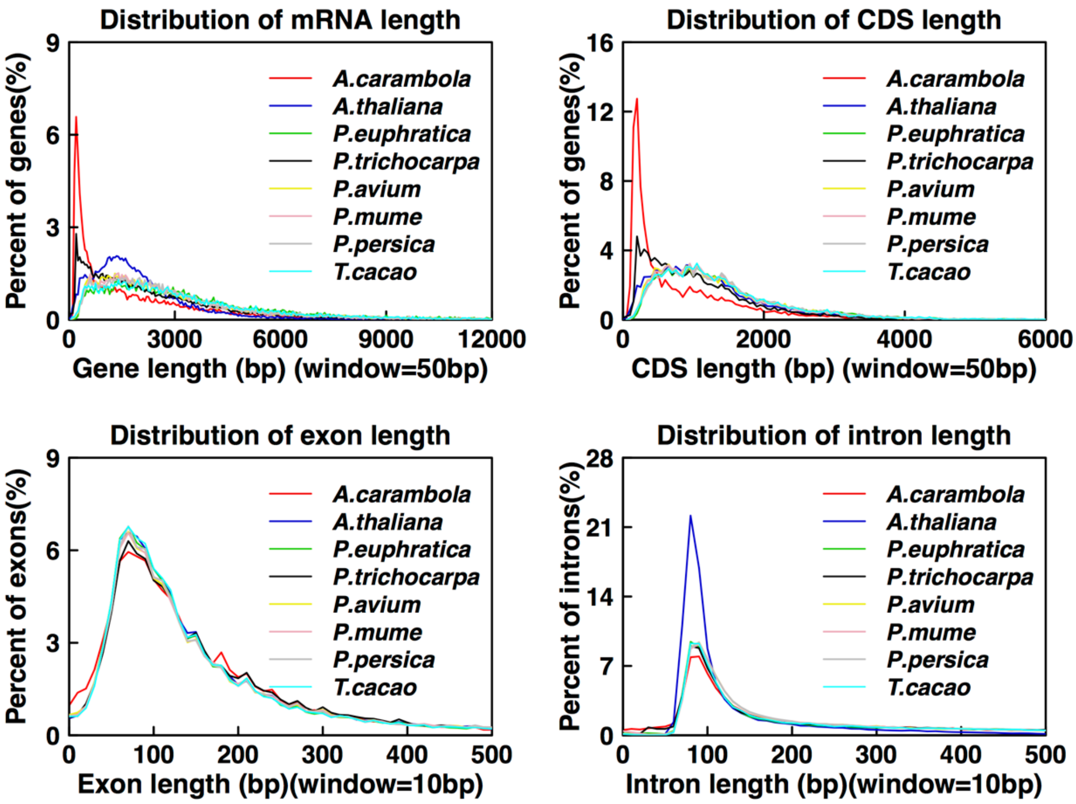


**Supplementary Figure 2. Comparison of the length distribution of gene sets in *A. carambola* and seven other plants**

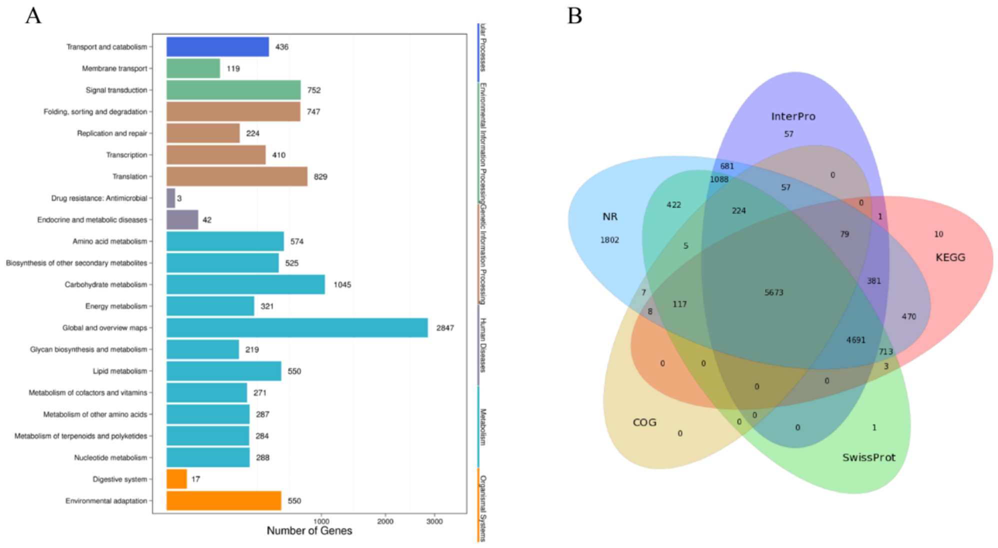


**Supplementary Figure 3. The statistics of gene annotation in *A. carambola.*** (A) The gene number distribution of KEGG pathway annotation; (B) The group of shared genes with five annotation methods.

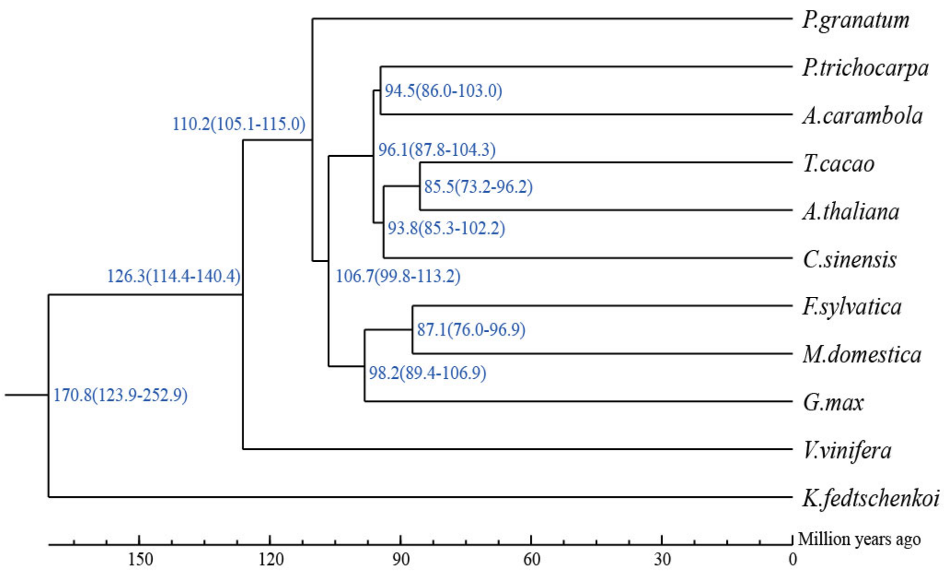


**Supplementary Figure 4. The phylogenetic tree with divergence time between *A. carambola* and other species were estimated using MCMCTREE with the default parameter.**
